# A versatile and stable reporter platform for human rhinovirus and other enteroviruses

**DOI:** 10.64898/2026.10.02.756346

**Authors:** Kazuma Okada, Junki Hirano, Takumi Adachi, Takafumi Noguchi, Kentaro Uemura, Chikako Ono, Shogo Nagata, Yuki Yamamoto, Tetsuharu Nagamoto, Shuhei Taguwa, Ming-Te Yeh, Yoshiharu Matsuura

**Affiliations:** Laboratory of Virus Control, Center for Infectious Disease Education and Research (CiDER), The University of Osaka, Osaka, Japan; Research Institute for Microbial Diseases (RIMD), The University of Osaka, Osaka, Japan; Center for Advanced Modalities and DDS (CAMaD), The University of Osaka, Osaka, Japan; HiLung Inc., Kyoto, Japan; Department of Microbiology and Immunology, National Cheng Kung University, Tainan, Taiwan.

## Abstract

Human rhinovirus (HRV), a major cause of upper respiratory infections, has increasingly been associated with lower respiratory tract disease. Although tools to investigate HRV pathogenesis are needed, they remain limited. Reporter viruses expressing fluorescent or luminescent proteins are powerful tools for analyzing viral distribution and replication; however, insertion of a reporter gene often impairs viral replication, and the inserted gene is readily lost during serial passage. To address these problems, we established a versatile and stable strategy for generating reporter HRV and other enteroviruses. The gene encoding mScarlet3, a red fluorescent protein, was inserted between the 5′UTR and VP4 gene, a site previously used in other enteroviruses but not in HRV. Optimizing the 2A cleavage sequence between mScarlet3 and VP4 yielded a reporter HRV with growth kinetics comparable to those of the wild-type virus. To improve the genetic stability of the reporter insert, we evaluated GC-content optimization of the reporter gene and relocation of the cis-acting replication element (CRE). GC-content optimization substantially improved reporter stability in both HRV-B14 and EV-A71, whereas CRE relocation provided additional stabilization in EV-A71 but not in HRV-B14, demonstrating the potential applicability of this strategy to other enteroviruses. This strategy for constructing stable reporter HRV and other enteroviruses will facilitate studies of enterovirus pathogenesis.

**IMPORTANCE:** Human rhinovirus (HRV) has lacked a reporter system expressing widely used reporter proteins, such as GFP, that enables visualization of viral dynamics in infected cells and tissues. The only reporter HRV reported to date loses fluorescence upon fixation and tolerates only small inserts, limiting its utility. Meanwhile, the clinical importance of HRV has become increasingly recognized, as reflected by its high detection rate among hospitalized children and recent outbreaks associated with fatal cases. A versatile and stable strategy for constructing reporter HRV is therefore urgently needed. Here, we overcome the two major obstacles to generating stable reporter HRV - impaired viral replication and loss of the inserted gene - and demonstrate that this strategy is also applicable to other enteroviruses. By providing a broadly applicable framework for constructing stable reporter viruses, this work establishes a foundation for visualizing infection dynamics and studying the pathogenesis of HRV and other enteroviruses.

## INTRODUCTION

Human rhinovirus (HRV) is a member of the genus *Enterovirus* within the family *Picornaviridae* and is the leading cause of the common cold. HRV exhibits high genetic diversity and is classified into three species: Enterovirus alpharhino (HRV-A), betarhino (HRV-B), and cerhino (HRV-C), comprising 80, 32, and 57 genotypes, respectively, with more than 160 genotypes identified to date (1). Although HRV has traditionally been regarded as a cause of mild upper respiratory illness, its clinical significance has become increasingly recognized in recent years. Accumulating evidence indicates that HRV can cause both upper and lower respiratory tract diseases, including pneumonia, asthma exacerbation, and chronic obstructive pulmonary disease (COPD) (1–4). In addition, several studies have reported that HRV is associated with higher rates of hospitalization and more frequent cardiopulmonary complications than influenza virus infection in both children and older adults (5, 6). Importantly, an outbreak of HRV-B14 in France in 2024 was associated with severe respiratory illness and multiple deaths (7). These observations underscore the growing public health importance of HRV and highlight the need for a better understanding of its pathogenesis. However, experimental tools available for studying HRV pathogenesis, including reporter viruses, remain limited.

Reporter viruses expressing fluorescent or luminescent proteins are powerful tools for studying viral infection and replication, enabling real-time visualization of viral spread, high-throughput screening of antiviral compounds and neutralizing antibodies, and non-invasive imaging of infection dynamics *in vivo*. However, for HRV, only a single reporter virus has been reported. Han *et al*. (8) generated a reporter HRV expressing the flavin-based (LOV-domain) fluorescent protein iLOV by inserting its 375-nt coding sequence between the VP1 and 2A regions. However, this approach faced two major limitations. First, this insertion site does not accommodate larger reporter genes, such as GFP or luciferase. Second, the recombinant virus exhibited attenuated growth compared with the wild-type virus, and iLOV fluorescence was lost upon fixation.

The construction of a stable reporter HRV presents several technical challenges. Reporter genes have most commonly been inserted at the VP1-2A junction, a strategy that requires identical protease cleavage sequences to flank the reporter gene and thereby promotes loss of the insert through homologous recombination (8). An alternative insertion site between the 5′UTR and VP4 has been used successfully in other enteroviruses (9). However, this approach requires the introduction of a new protease cleavage sequence between the reporter and VP4, and systematic optimization of this sequence to ensure efficient reporter expression while maintaining viral replication has received little attention. Furthermore, reporter gene instability is a common problem among enteroviruses, in which larger inserts are frequently lost during serial passage or constrained by genome packaging capacity (10, 11). Therefore, no genetically stable reporter HRV carrying a large reporter gene has yet been developed.

In this study, we established a strategy for generating versatile and genetically stable reporter HRVs. Using an efficient plasmid-based recovery system, we inserted the full-length coding sequence of mScarlet3, a red fluorescent protein, between the 5′UTR and VP4 gene of the HRV genome. Although this region has been used for reporter insertion in other enteroviruses (9), its application to HRV has not previously been reported. To overcome reporter gene instability, we employed a two-pronged strategy combining GC-content optimization of the inserted gene with relocation of the *cis*-acting replication element (CRE). We first established this strategy in HRV-B14 and subsequently applied it to enterovirus A71 (EV-A71) to evaluate its broader applicability across other enteroviruses.

## RESULTS

### Identification of the optimal plasmid vector for recombinant rhinovirus recovery

To identify the most efficient system for recovering recombinant HRV, we constructed four plasmid vectors. We used two transcription promoters, RNA polymerase I (Pol I) and an RNA polymerase II promoter (CMV), that had been previously evaluated for recombinant virus recovery using an internal ribosome entry site (IRES) (12). For each promoter, an additional construct containing a hammerhead ribozyme sequence upstream of the HRV genome was generated. The full-length genomes of HRV-A1B and HRV-B14 were cloned into the resulting four plasmids: Pol I, Pol I with an upstream hammerhead ribozyme (P+hh), CMV, and CMV with an upstream hammerhead ribozyme (C+hh) (Fig. 1A). Each plasmid was transfected into H1-HeLa cells, and viral titers in the culture supernatant were determined by plaque assay at 4 days post-transfection (dpt). For both HRV-A1B and HRV-B14, the Pol I vector consistently yielded viral titers approximately 100-fold higher than those obtained with the other vectors (Fig. 1B and 1C).

**FIG 1.**
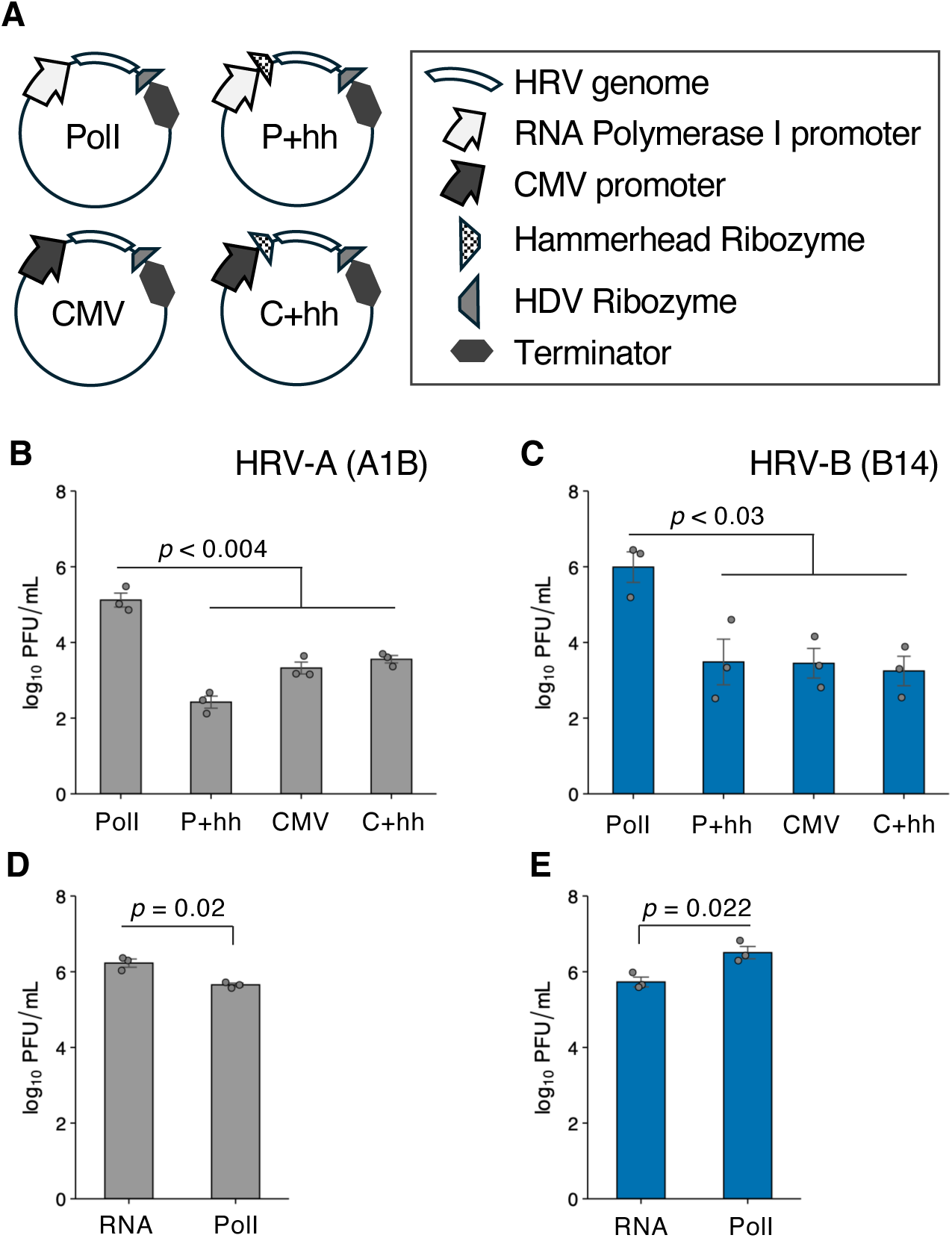
RNA polymerase I–driven plasmids enable efficient recovery of recombinant HRV. (A) Schematic of the four reverse genetics vectors used to recover recombinant HRV. Each construct carries the HRV genome under the control of the RNA polymerase I (Pol I) or CMV promoter, with or without an upstream hammerhead ribozyme, followed by an HDV ribozyme and a transcription terminator. (B, C) Recombinant virus recovery from the four vector types for HRV-A1B (B) and HRV-B14 (C). The indicated plasmids were transfected into H1-HeLa cells, and viral titers in the culture supernatant were determined by plaque assay at 4 dpt. (D, E) Comparison of viral recovery between Pol I plasmid transfection and RNA transfection for HRV-A1B (D) and HRV-B14 (E), determined by plaque assay at 4 dpt. Data are presented as mean ± SD from three independent biological replicates. *P* values were determined by Welch’s *t*-test with Holm correction (B, C) or Welch’s *t*-test (D, E); exact values are indicated in each panel.

Because the Pol I vector showed the highest recovery efficiency, we next compared viral recovery following Pol I plasmid transfection with that following *in vitro*-transcribed RNA transfection. For HRV-A1B, RNA transfection produced higher viral titers than Pol I plasmid transfection (Fig. 1D). In contrast, for HRV-B14, Pol I plasmid transfection yielded higher titers than RNA transfection (Fig. 1E). Overall, the Pol I plasmid-based recovery system was comparable to, and in some strains more efficient than, the conventional RNA transfection method.

### Selection of optimal amino acid sequences for 2A- or 3C-mediated cleavage between mScarlet3 and VP4

Building on the plasmid-based recovery system for HRV-B14, we next generated a reporter B14 virus. We selected the region between the 5′UTR and VP4 gene as the insertion site for the reporter gene encoding mScarlet3, a red fluorescent protein (Fig. 2A) (13). Because mScarlet3 is translated as part of the viral polyprotein and fused to VP4, efficient proteolytic cleavage between mScarlet3 and VP4 was expected to be critical for both reporter expression and viral replication (Fig. 2A).

**FIG 2.**
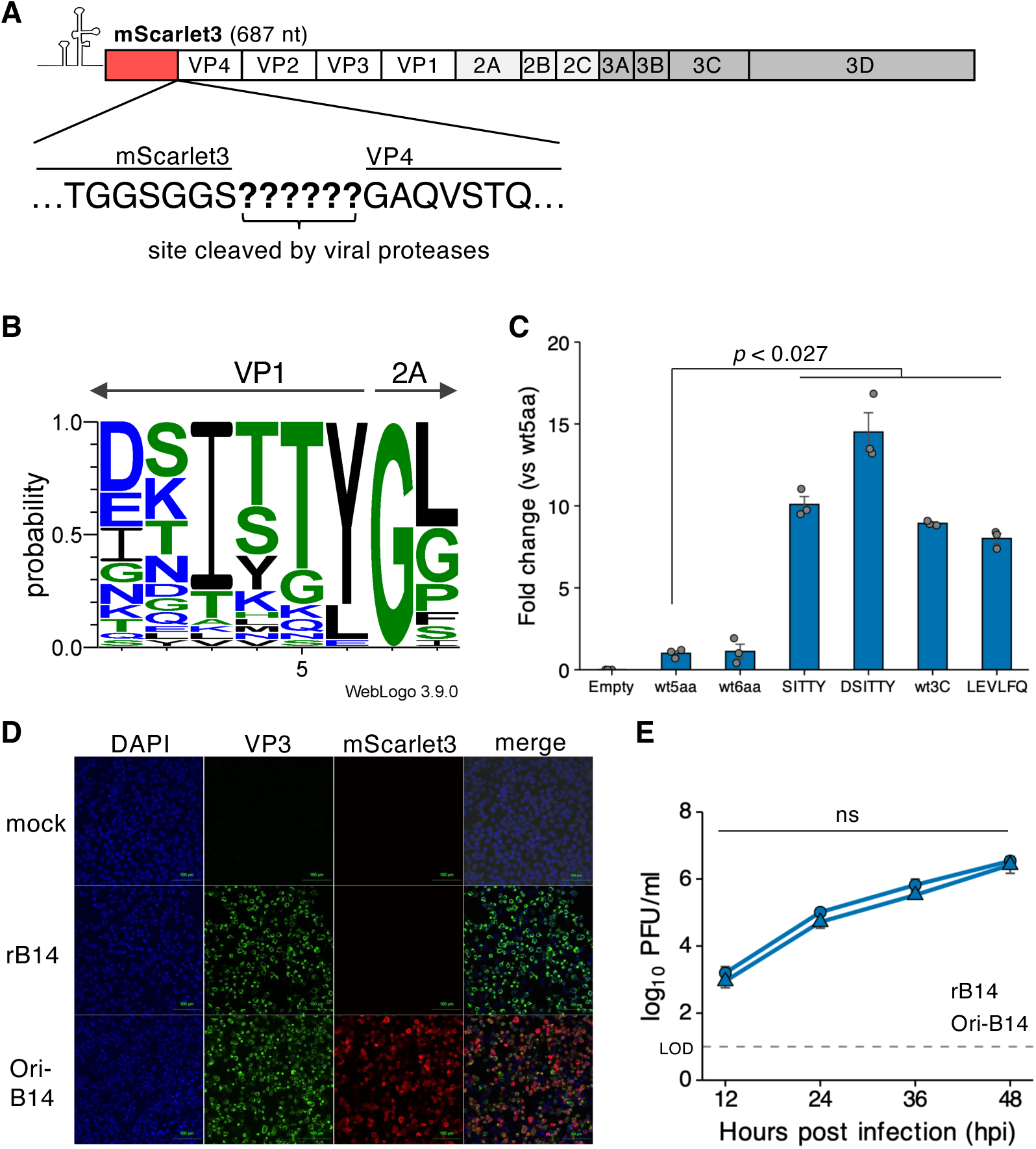
Optimization of the 2A cleavage site enables reporter expression while preserving HRV replication. (A) Schematic of the recombinant HRV genome expressing mScarlet3. The 687-nt mScarlet3 gene (red) was inserted between the 5′UTR and VP4 of HRV-B14, with a protease cleavage sequence placed between mScarlet3 and VP4 so that mScarlet3 is released from the viral polyprotein by viral proteases. (B) Alignment of the VP1–2A junction sequences of 32 HRV-B strains (Table S1), generated with MAFFT and visualized using WebLogo. (C) Genome plasmids encoding mScarlet3 with each of six cleavage sequences (wt5aa, wt6aa, SITTY, DSITTY, wt3C, or LEVLFQ) between mScarlet3 and VP4, or an empty vector lacking mScarlet3 (Empty), were transfected into H1-HeLa cells, and the proportion of red-fluorescent cells was determined by flow cytometry at 4 dpt. Each construct was compared with wt5aa. (D) H1-HeLa cells were infected with Ori-B14, which carries mScarlet3 with the DSITTY cleavage sequence, at an MOI of 0.1, fixed at 12 hpi, and stained with an anti-VP3 antibody and DAPI. mScarlet3 (red) was detected by its intrinsic fluorescence. Mock-, rB14-, and Ori-B14–infected cells are shown. Scale bar, 100 µm. (E) Growth kinetics of rB14 and Ori-B14. H1-HeLa cells were infected at an MOI of 0.01, and viral titers in the culture supernatant were determined by plaque assay at 12, 24, 36, and 48 hpi. Data are presented as mean ± SD from three independent biological replicates. *P* values were determined by Welch’s *t*-test with Holm correction (C, E); exact values are indicated in each panel.

To identify an optimal cleavage sequence, we aligned the 2A protease cleavage sequences spanning the VP1-2A junction from representative HRV-B genotypes using MAFFT (Fig. 2B, Table S1). The 5-amino-acid sequence SITTY and the 6-amino-acid sequence DSITTY were the most highly conserved motifs. In addition to these consensus sequences, we constructed HRV-B14 genomic plasmids carrying the native B14 5-amino-acid (wt5aa) and 6-amino-acid (wt6aa) 2A cleavage sequences between mScarlet3 and VP4. For comparison, we also generated constructs containing either a commercially available 3C protease cleavage sequence (LEVLFQ), previously shown to be efficiently cleaved by the HRV-B14 3C protease, or the native B14 3C cleavage sequence (wt3C), at the same position.

These plasmids were transfected into H1-HeLa cells, and cells were harvested at 4 dpt for flow cytometric analysis of mScarlet3-positive cells. Using the native B14 cleavage sequence (wt5aa) as the reference, DSITTY yielded the highest proportion of fluorescent cells among the constructs that showed significantly higher reporter expression (Fig. 2C, Fig. S1). We therefore selected the reporter virus carrying the DSITTY cleavage sequence, designated Ori-B14, for subsequent analyses. To confirm reporter expression, H1-HeLa cells infected with Ori-B14 were immunostained with an anti-VP3 antibody at 12 hours post-infection (hpi). mScarlet3 fluorescence (red) co-localized with VP3-positive cells (green), confirming reporter expression in the infected cells (Fig. 2D). The growth kinetics of Ori-B14 were comparable to those of the non-reporter recombinant virus (rB14), with no significant differences observed at any time point (Fig. 2E). These results indicate that optimization of the protease cleavage sequence between the reporter protein and VP4 enables efficient reporter expression while preserving viral replication.

### Genetic stability of the mScarlet3 gene inserted between the 5′UTR and VP4 in HRV-B14

Because previous studies showed that reporter inserts longer than GFP are unstable in reporter HRV (8), we next examined the stability of the mScarlet3 gene in Ori-B14. We infected H1-HeLa cells with Ori-B14 at a multiplicity of infection (MOI) of 0.01 and virus was serially passaged five times under the same conditions. Viruses from the first (P1) and fifth (P5) passages were inoculated into H1-HeLa at an MOI of 0.1 and cells were stained with an anti-VP3 antibody at 12 hpi and analyzed by flow cytometry for VP3 and mScarlet3 expression. VP3-positive, mScarlet3-negative cells, indicative of infection by viruses lacking the reporter gene, were rarely detected in P1 but became abundant in P5 (Fig. 3A). Consistent with this observation, the mScarlet3 retention, defined as the proportion of mScarlet3-positive cells among VP3-positive cells, decreased significantly from 94% in P1 to 14% in P5 (Fig. 3B). RT-PCR analysis of viral RNA detected a shorter-than-expected band only in P5 (Fig. 3C), and sequencing of this band revealed a deletion spanning 197-622 nt of the mScarlet3 open reading frame (ORF) (Fig. 3D).

**FIG 3.**
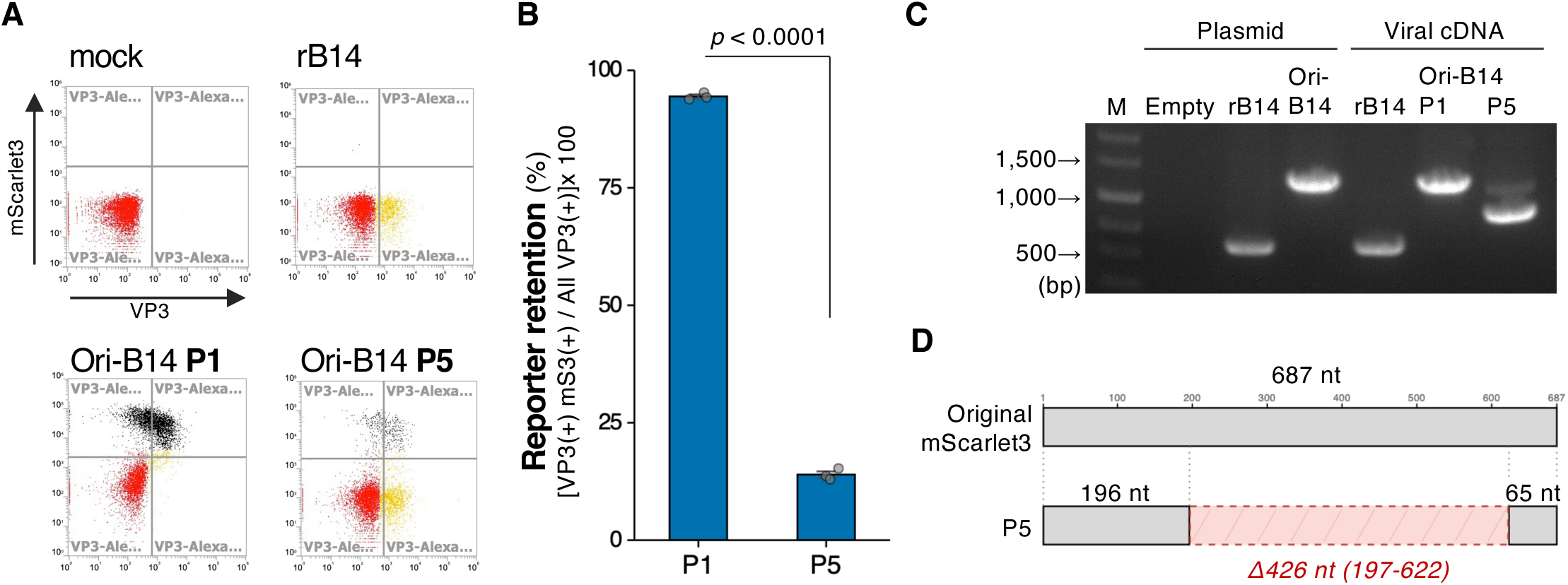
The mScarlet3 insert is lost during serial passage of Ori-B14. Ori-B14 was serially passaged in H1-HeLa cells at an MOI of 0.01, and the first (P1) and fifth (P5) passages were analyzed. (A) H1-HeLa cells were infected with P1 or P5 at an MOI of 0.1, fixed at 12 hpi, and stained with an anti-VP3 antibody. mScarlet3 (y axis) and VP3 (x axis) signals were analyzed by flow cytometry. Mock- and rB14-infected cells served as controls. VP3-positive, mScarlet3-negative cells (lower right) indicate infection by a mScarlet3-deficient mutant. (B) Reporter retention, calculated as the proportion of mScarlet3-positive cells among VP3-positive cells [VP3(+), mScarlet3(+) / all VP3(+) × 100], for P1 and P5. (C) RT-PCR of the region spanning the 5′UTR to the 5′ end of VP4. RNA extracted from P1 and P5 was reverse-transcribed and amplified. The corresponding genome plasmids (Empty, rB14, and Ori-B14) and viral cDNA from rB14 were included as references. M, size marker. (D) Schematic of the deletion identified by sequencing the shorter P5 amplicon. Relative to the 687-nt mScarlet3 ORF, a 426-nt deletion (nt 197–622) was detected, leaving 196 nt at the 5′ end and 65 nt at the 3′ end. Data are presented as mean ± SD from three independent biological replicates. The *P* value was determined by Welch’s *t*-test and is indicated in the panel.

These results indicated that optimization of the protease cleavage sequence alone is insufficient to maintain genetic stability of the inserted gene, highlighting the need for additional strategies to prevent reporter gene loss.

### Improved genetic stability of the mScarlet3 gene inserted in HRV-B14 and in EV-A71 through GC content optimization

To improve the genetic stability of the reporter insert in HRV, we investigated the effect of GC content. We first analyzed GC content across the HRV-B14 genome using a 100-nucleotide sliding window with a 50-nucleotide step size. The region between the 5′UTR and VP4 gene exhibited the greatest variation in GC content (Fig. 4A). Because mScarlet3 was inserted into this region, we hypothesized that adjusting the GC content of the reporter gene to more closely match that of the surrounding viral genome would improve its genetic stability.

**FIG 4.**
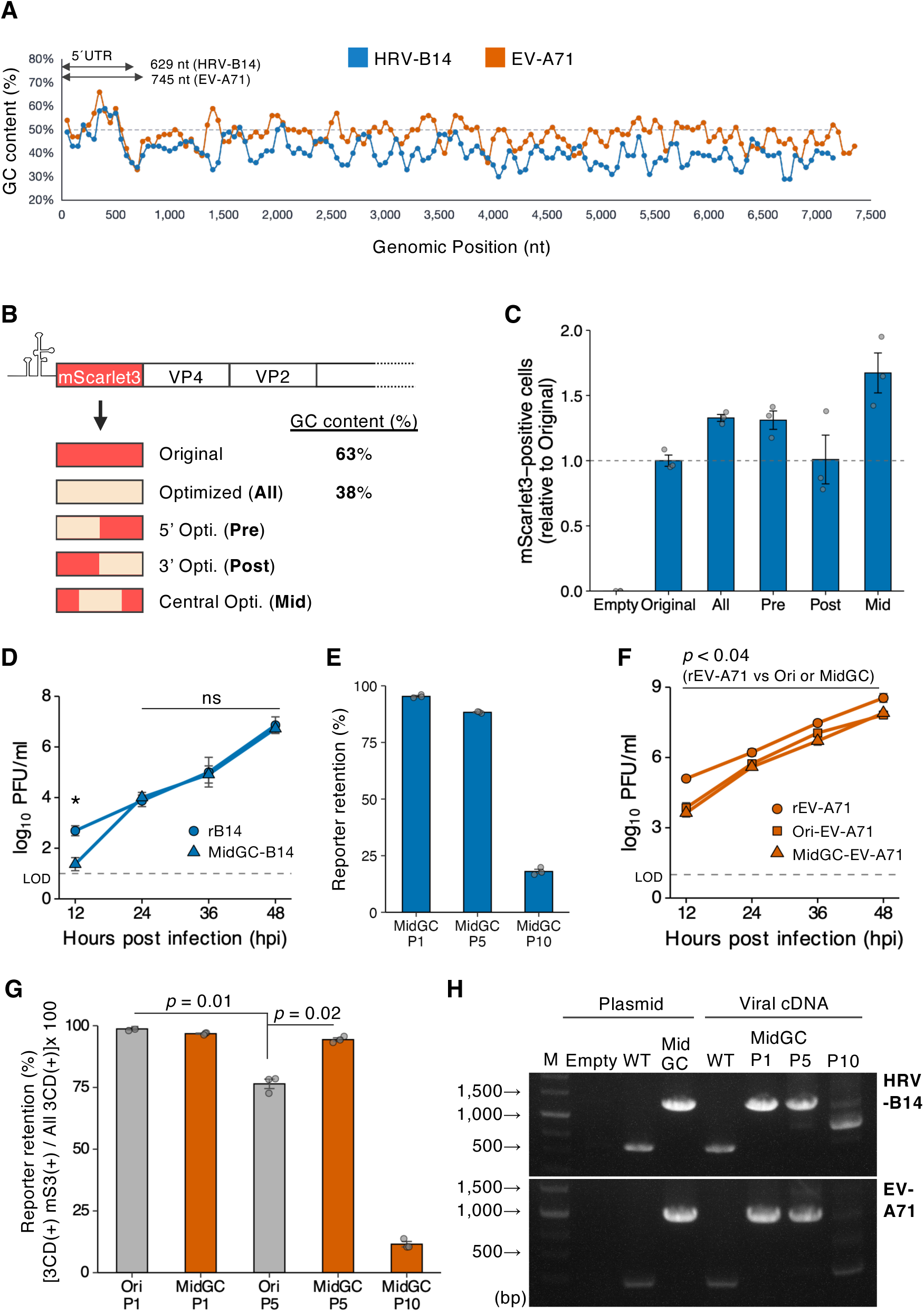
GC-content optimization of the insert improves stability in HRV-B14 and EV-A71. (A) GC content across the HRV-B14 (blue) and EV-A71 (vermilion) genomes, calculated using a 100-nt sliding window with a 50-nt step. The shaded area marks the 5′UTR–VP4 region, which showed the greatest variation in GC content. (B) Schematic of the mScarlet3 GC-content variants. The native mScarlet3 (Original, 63% GC) was modified to reduce the GC content to 38% across the entire gene (All) or within the 5′ (Pre), 3′ (Post), or central (Mid) region. (C) Genome plasmids carrying each mScarlet3 variant, or an empty vector lacking mScarlet3 (Empty), were transfected into H1-HeLa cells, and the proportion of mScarlet3-positive cells, normalized to that of Original, was determined by flow cytometry at 4 dpt. (D) Growth kinetics of rB14 and MidGC-B14. H1-HeLa cells were infected at an MOI of 0.01, and viral titers in the culture supernatant were determined by plaque assay at 12, 24, 36, and 48 hpi. (E) Reporter retention of MidGC-B14 at P1, P5, and P10. (F) Growth kinetics of rEV-A71, Ori-EV-A71, and MidGC-EV-A71. RD-S cells were infected at an MOI of 0.01, and viral titers were determined by plaque assay at 12, 24, 36, and 48 hpi. (G) Reporter retention of Ori-EV-A71 and MidGC-EV-A71 at the indicated passages, calculated as the proportion of mScarlet3-positive cells among 3CD-positive cells [3CD(+), mScarlet3(+) / all 3CD(+) × 100]. (H) RT-PCR of the region spanning the 5′UTR to the 5′ end of VP4 for HRV-B14 (top) and EV-A71 (bottom). RNA from the indicated passages was reverse-transcribed and amplified. The corresponding genome plasmids (Empty, WT, and MidGC) and viral cDNA from WT were included as references; WT refers to rB14 (HRV-B14) or rEV-A71 (EV-A71). M, size marker. Data are presented as mean ± SD from three independent biological replicates. The limit of detection (LOD) is indicated by a dashed line in D and F. Statistical comparisons used the Games-Howell test (C, E, G), with no significant differences detected in C; a per-time-point Welch’s *t*-test with Holm correction (D); or Welch’s *t*-test with Holm correction against rEV-A71 (F). *P* values are indicated in each panel. ns, not significant.

The native GC content of the mScarlet3 coding sequence is 63%, near the upper end of the range observed across the HRV-B14 genome (Fig. 4A). We therefore generated a variant in which the GC content of the entire mScarlet3 coding sequence was reduced to 38% (designated All), approximating the average GC content of the HRV-B14 genome (Fig. 4B). To determine whether local rather than global optimization was sufficient, we also generated variants in which only the 5’ region (Pre), the central region (Mid), or the 3’ region (Post) was reduced to 38% GC (Fig. 4B).

Genomic plasmids carrying each GC-optimized mScarlet3 variant together with the DSITTY cleavage sequence were transfected into H1-HeLa cells, and reporter expression was evaluated by flow cytometry at 4 dpt. All GC-optimized variants exhibited proportions of mScarlet3-positive cells comparable to or higher than that of Ori-B14, which carries the original, non-optimized mScarlet3 sequence (Fig. 4C), indicating that GC-content optimization did not impair reporter expression. Because the deletion detected in Ori-B14 after serial passage was located within the central region of mScarlet3 coding sequence (Fig. 3D), and because the Mid variant consistently showed the highest level of reporter expression, we selected the GC-optimized Mid variant (MidGC-B14) for subsequent analyses.

The growth kinetics of MidGC-B14 were comparable to those of rB14, although viral titers were significantly lower at 12 hpi (Fig. 4D). We next assessed reporter stability by serial passage. Whereas the reporter retention of Ori-B14 declined markedly by P5, MidGC-B14 retained the reporter gene in 88% of infected cells at P5, although the retention decreased to 18% by P10 (Fig. 4E, Fig. S2). RT-PCR analysis of RNAs from P1 to P10 detected a shorter-than-expected band only in P10 (Fig. 4H). These findings demonstrated that GC-content optimization substantially improves the genetic stability of the inserted reporter gene in HRV-B14.

To determine whether GC-content optimization also stabilizes reporter inserts in enteroviruses other than HRV, we applied the same strategy to EV-A71. Sliding-window analysis of the EV-A71 genome revealed that, as in HRV-B14, the region between the 5′UTR and VP4 gene exhibited the greatest variation in GC content (Fig. 4A).

We next compared the growth kinetics of reporter EV-A71 carrying either the original mScarlet3 sequence (Ori-EV-A71) or the GC-optimized Mid variant (MidGC-EV-A71) with those of the non-reporter recombinant virus (rEV-A71). Although both reporter viruses showed significantly lower titers than rEV-A71 at all time points, they reached nearly 10^8^ PFU/ml by 48 hpi (Fig. 4F). Because the protease cleavage sequence between the reporter gene and VP4 was adopted from a previous study (14), further optimization of this sequence may improve the replication of reporter EV-A71.

To assess insert stability, Ori-EV-A71 and MidGC-EV-A71 were serially passaged in RD-S cells at an MOI of 0.01 to generate the first (P1), fifth (P5), and tenth (P10) passages. Cells infected with P1, P5, or P10 at an MOI of 0.1 were harvested at 14 hpi, when cytopathic effect became apparent, stained with an anti-3CD antibody, and analyzed by flow cytometry. Reporter retention was calculated as the proportion of mScarlet3-positive cells among all 3CD-positive cells, as described for HRV-B14. At P5, MidGC-EV-A71 exhibited significantly higher reporter retention than Ori-EV-A71 (94.4% versus 76.5%; Fig. 4G, Fig. S2). Moreover, reporter retention in MidGC-EV-A71 remained unchanged between P1 and P5 (96.8% and 94.4%), whereas Ori-EV-A71 showed significant decline over the same period. However, reporter retention in MidGC-EV-A71 decreased to 11.5% by P10 (Fig. 4G). RT-PCR analysis of RNAs from P1 to P10 showed a shorter-than-expected band in P10 (Fig. 4H).

These findings demonstrate that GC-content optimization substantially improves reporter gene stability in both HRV-B14 and EV-A71 but is insufficient to maintain long-term genetic stability, indicating that additional strategies are required.

### Improved genetic stability of the mScarlet3 insert through CRE relocation

To further enhance reporter gene stability, we next employed relocation of the *cis*-acting replication element (CRE). The CRE is an internal RNA structure required for genome replication. It was first identified in HRV-B14 (15, 16) and subsequently in poliovirus and other picornaviruses, where it can retain its function after relocation to an alternative genomic position (17). CRE relocation has previously been used to stabilize genetically modified poliovirus vaccine strains (18, 19). We therefore investigated whether this strategy could also improve the stability of the inserted reporter gene.

We first sought to identify a region of mScarlet3 that could accommodate the CRE without impairing fluorescence. Based on structural predictions generated by AlphaFold2, three candidate insertion sites predicted to preserve the structure required for fluorescence were selected (Fig. S3A and S3B). H1-HeLa cells were transfected with plasmids encoding mScarlet3 containing the HRV-B14 CRE inserted at one of three sites (75CRE, 186CRE, or 229CRE), or with the original mScarlet3 construct, and reporter expression was determined by flow cytometry at 24 hpt. The 75CRE and 186CRE constructs exhibited significantly reduced fluorescence compared with the original mScarlet3, whereas 229CRE construct showed comparable fluorescence (Fig. S3C). We therefore selected the 229CRE construct for subsequent experiments.

Previous studies on poliovirus vaccine development showed that the relocated CRE functions efficiently only after inactivation of the native CRE by mutation (18, 19). Based on the MidGC reporter viruses, we therefore generated, in both HRV-B14 and EV-A71, three additional constructs carrying a deficient mutation in CRE (ΔCRE), a relocation of CRE (Mid_CRE), or both modifications (Mid_CRE/ΔCRE) (Fig. 5A). These genomic plasmids were transfected into H1-HeLa cells (HRV-B14) or RD-S cells (EV-A71), and red fluorescence was assessed at 4 dpt. In both viruses, the ΔCRE construct produced almost no fluorescence, consistent with defective viral replication (Fig. 5B). In contrast, both Mid_CRE and Mid_CRE/ΔCRE exhibited robust fluorescence indicative of viral replication. These viruses were therefore used for subsequent analyses of replication and reporter gene stability.

**FIG 5.**
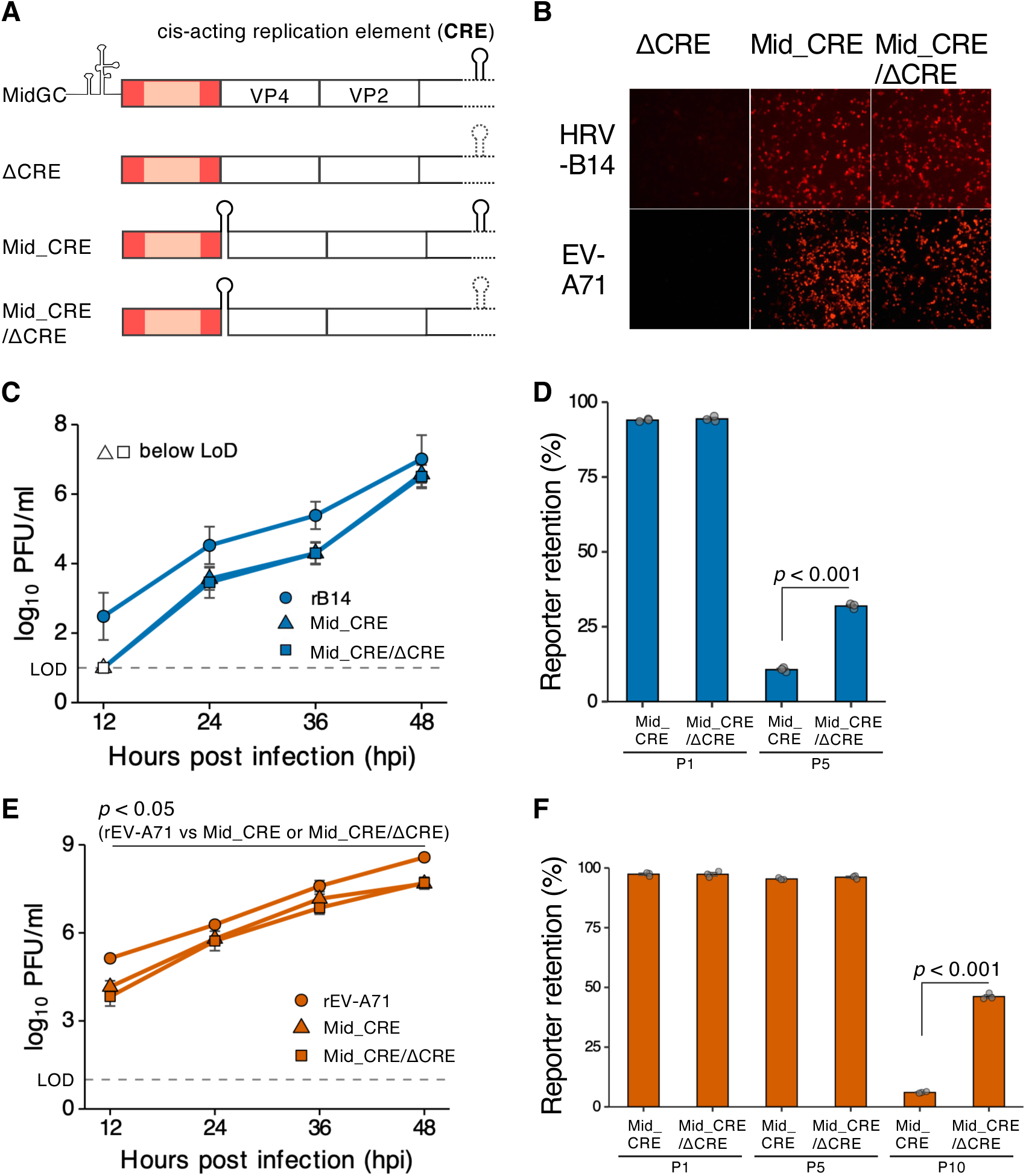
CRE relocation stabilizes the insert in EV-A71 but not in HRV-B14. (A) Schematic of the constructs used to test CRE relocation. Starting from the MidGC strain, strains were generated carrying an inactivating mutation in the native CRE (ΔCRE), a CRE relocated into mScarlet3 (Mid_CRE), or both (Mid_CRE/ΔCRE). Inactivated CREs are shown as dashed stem-loops. (B) Genome plasmids of each construct were transfected into H1-HeLa (HRV-B14) or RD-S (EV-A71) cells, and mScarlet3 fluorescence was imaged at 4 dpt. (C) Growth kinetics of rB14, Mid_CRE, and Mid_CRE/ΔCRE of HRV-B14. H1-HeLa cells were infected at an MOI of 0.01, and viral titers in the culture supernatant were determined by plaque assay at 12, 24, 36, and 48 hpi. Data points below the limit of detection (LOD) are shown as open symbols. (D) Reporter retention of Mid_CRE and Mid_CRE/ΔCRE of HRV-B14 at P1 and P5. (E) Growth kinetics of rEV-A71, Mid_CRE, and Mid_CRE/ΔCRE of EV-A71. RD-S cells were infected at an MOI of 0.01, and viral titers were determined by plaque assay at 12, 24, 36, and 48 hpi. (F) Reporter retention of Mid_CRE and Mid_CRE/ΔCRE of EV-A71 at P1, P5, and P10. Data are presented as mean ± SD from three independent biological replicates. The LOD is indicated by a dashed line in C and E. Statistical comparisons used Welch’s *t*-test with Holm correction against rB14 (C), or against rEV-A71 (E), or the Games-Howell test (D, F); *P* values are indicated in each panel.

For HRV-B14, the titers of both Mid_CRE and Mid_CRE/ΔCRE approached those of rB14 by 48 hpi, although they remained approximately 1 log10 PFU/ml lower between 12 and 36 hpi (Fig. 5C). We next assessed reporter gene stability by serial passage. Whereas MidGC retained approximately 90% of the reporter gene expression at P5 (Fig. 4E, Fig. S2), reporter retention decreased to 10.7% and 31.9% in Mid_CRE and Mid_CRE/ΔCRE, respectively (Fig. 5D, Fig. S4A). Thus, CRE relocation did not improve reporter gene stability in HRV-B14; instead, it reduced stability relative to MidGC, indicating that GC-content optimization alone was the most effective strategy in HRV-B14.

For EV-A71, the titers of both Mid_CRE and Mid_CRE/ΔCRE remained significantly lower than those of rEV-A71 at all time points but reached approximately 10^8^ PFU/ml by 48 hpi (Fig. 5E). Both Mid_CRE and Mid_CRE/ΔCRE retained nearly 100% of the reporter at P5 (Fig. 5F, Fig. S4B). At P10, reporter retention in MidGC-EV-A71 had declined to approximately 10% but retained 46.2% in Mid_CRE/ΔCRE, indicating that CRE relocation improved genetic stability in EV-A71. In contrast, reporter retention in Mid_CRE decreased to 6.1% at P10, indicating that stabilization of the reporter gene required both relocation of the CRE and inactivation of the original CRE (Fig. 5F).

Finally, to explore whether this strategy might be applicable beyond HRV-B and EV-A, we used RNA secondary-structure prediction to examine whether the relocated CRE retained its structure in other enterovirus species (Fig. 6). Seven viruses were analyzed: HRV-A16, HRV-B14, HRV-C15, EV-A71, coxsackievirus B3 (CVB3), Poliovirus type1 (PV1), and EV-D68. For each virus, we modeled the junction formed by the 3′-terminal 100 nt of the Mid variant, the relocated CRE, and the 5′-terminal 100 nt of the VP4 gene.

**FIG 6.**
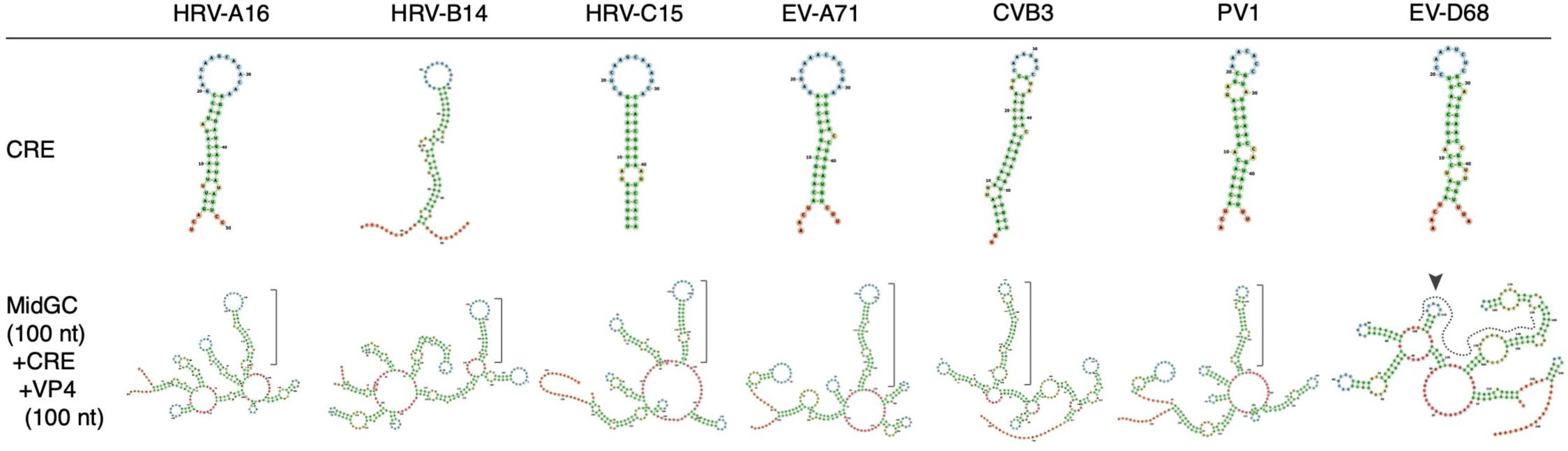
The predicted secondary structure of the relocated CRE varies among enterovirus species. RNA secondary structures were predicted for the CRE region of seven enterovirus and rhinovirus strains (HRV-A16, HRV-B14, HRV-C15, EV-A71, CVB3, PV1, and EV-D68). Top, the predicted structure of the CRE alone. Bottom, the predicted structure of the junction region modeled in the MidGC reporter context, comprising the relocated CRE and its flanking sequences. Brackets indicate the maintained CRE stem-loop. The critical CRE structure was predicted to be maintained in HRV-A16, HRV-B14, HRV-C15, EV-A71, CVB3, and PV1, but disrupted in EV-D68 (arrowhead).

The characteristic CRE secondary structure was predicted to be preserved in HRV-A16, HRV-B14, HRV-C15, EV-A71, CVB3, and PV1, but not in EV-D68. These findings suggest that although CRE relocation is broadly applicable across enteroviruses, additional sequence optimization may be required in some species to maintain the functional CRE structure.

### MidGC-B14 combined with airway organoids can recapitulate infection dynamics in the airway epithelium

To evaluate the utility of the reporter HRV, we inoculated MidGC-B14 into human induced pluripotent stem cell (iPSC)-derived airway organoids (20, 21) and monitored viral propagation by the expression of red fluorescence together with the antiviral immune response (Fig. 7). Because the organoids were maintained in an air-liquid interface system, the basolateral medium was collected over time to measure host factors involved in the immune response.

**FIG 7.**
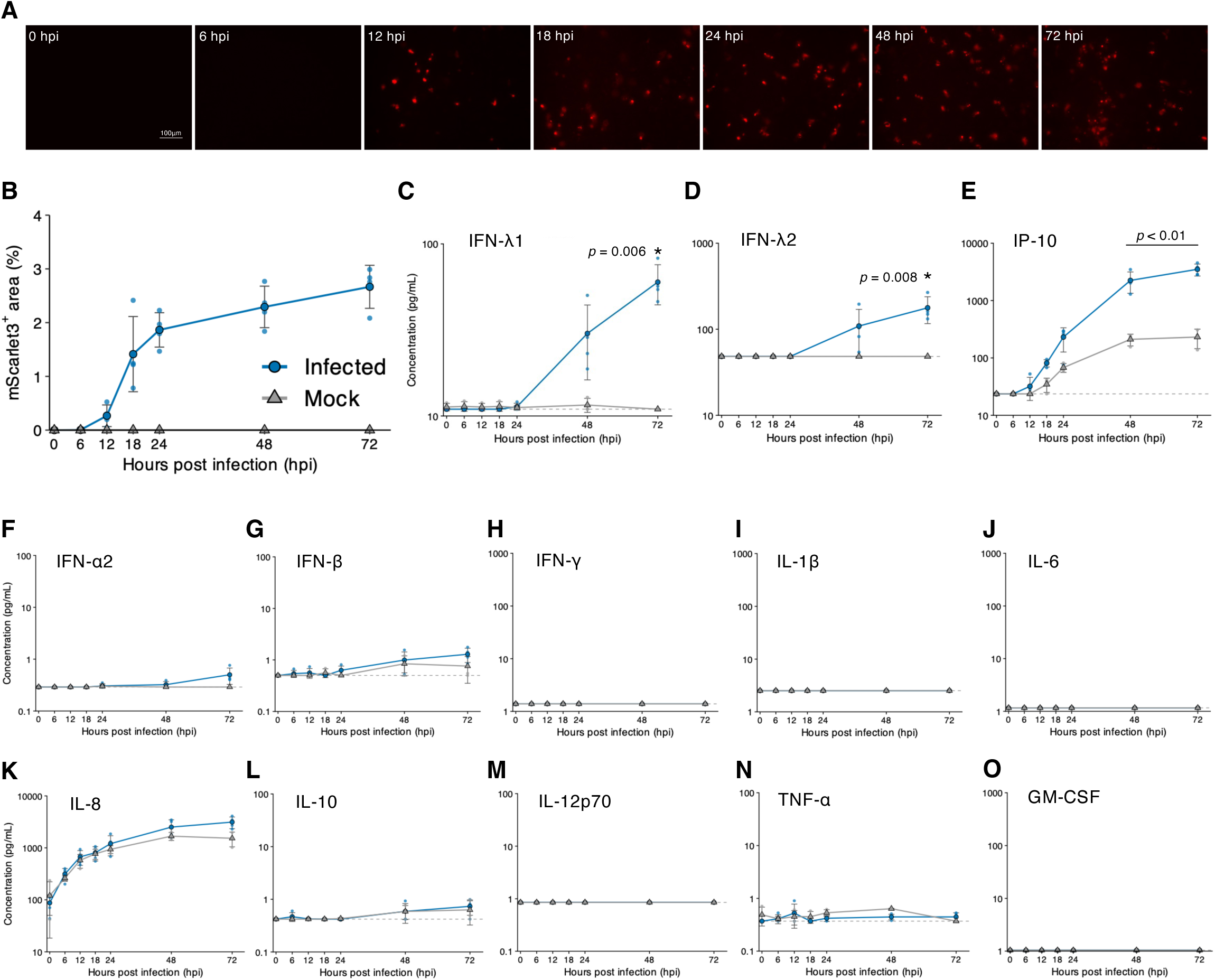
MidGC-B14 combined with airway organoids recapitulates infection dynamics in the airway epithelium. (A) The airway epithelial organoid was infected with MidGC-B14 at an MOI of 0.1 and imaged at 0, 6, 12, 18, 24, 48, and 72 hpi. Representative images are shown. (B) The red fluorescence area was quantified at each time point. (C to O) Concentrations of 13 interferons and cytokines in the basolateral medium were measured at each time point. Blue circles indicate the infected group and gray triangles the mock group. The dashed line indicates the limit of detection (LOD). Values below the LOD were substituted with the LOD value for analysis and are plotted. Data are presented as mean ± SD (mock, n = 3; infected, n = 4). *P* values were determined by Welch’s *t*-test with Holm correction against the mock group and are indicated in each panel.

At 6, 12, 18, 24, 48, and 72 hpi, the red fluorescent area gradually expanded, indicating viral propagation (Fig. 7A and 7B). Quantification of interferons and cytokines in the medium collected at these time points showed that secretion of IFN-*λ*1 and IFN-*λ*2, key mediators of the airway mucosal immune response, significantly increased in infected organoids at 72 hpi (Fig. 7C and 7D). Among the 13 factors measured, IP-10 / CXCL10, a biomarker associated with HRV-induced asthma exacerbations (22), was the only factor that reached significance at an earlier time point, being elevated in infected organoids from 48 hpi (Fig. 7E). IFN-*α*2 and IFN-*β* were quantifiable from 24 and 6 hpi, respectively, and were slightly higher in infected organoids at 72 hpi, but neither difference was significant (Fig. 7F and G). IL-8 was quantifiable at all time points and increased over time in both groups, with no significant difference between the two groups at any time point (Fig. 7K). IL-10 and TNF-*α* were also quantifiable but did not differ significantly between the groups at any time point (Fig. 7L and N). IFN-*γ*, IL-1*β*, IL-6, IL-12p70, and GM-CSF were below the limit of detection (LOD) in all replicates of both groups at every time point (Fig. 7H-J, M, and O).

These results showed that combining MidGC-B14 with airway organoids enables real-time monitoring of viral spread through red fluorescence while simultaneously assessing antiviral immune responses in the airway epithelium. Moreover, the increased secretion of IFN-*λ*1, IFN-*λ*2, and IP-10 suggests that this system recapitulates key features of the airway epithelium immune response to HRV infection.

## DISCUSSION

Reporter enteroviruses, particularly HRVs, have been difficult to construct because foreign inserts are rapidly lost during viral replication (8). In this study, we addressed this limitation through a modular strategy consisting of reporter insertion at the 5′UTR–VP4 junction, optimization of the viral protease cleavage site, and stabilization of the insert by GC optimization and/or CRE relocation. These modifications generated stable, replication-competent reporter viruses in two representative enteroviruses, HRV-B14 and EV-A71. The successful application of the same strategy to both viruses suggests that it captures general principles governing insert stability across the Enterovirus genus.

The GC content of a gene strongly influences the stability of its RNA secondary structure, and both the GC content and sequence distribution of foreign inserts have been associated with their genetic stability in recombinant polioviruses, independently of insert size (23). Likewise, matching the codon usage and GC content of a foreign insert to those of the viral genome, rather than those of the host, has been reported to improve inserted gene stability in RNA virus vectors (24). In Ori-B14, the reporter gene retained the native GC content of mScarlet3 (63%), which is higher than the average GC content of the viral genome. Viral RdRp is thought to stall at stable RNA secondary structures and undergo template switching, deleting the intervening region (25, 26). Lowering the GC content of the insert is expected to reduce the propensity of the region to form such stable structures, thereby reducing the likelihood of template switching and improving reporter gene stability. Consistent with this, GC optimization of the central region, where the deletion occurred in Ori-B14, improved reporter stability, suggesting that reducing the local structural stability of the inserted reporter gene contributed to its genetic stability.

Insert stabilization by CRE relocation required inactivation of the native CRE. In the Mid_CRE virus, in which the native CRE remained active, the insert stability was comparable to that of the MidGC virus. In contrast, improved reporter stability was observed only in the Mid_CRE/ΔCRE virus, in which the native CRE was inactivated (Fig. 5F). We attribute this difference to the essential role of the relocated CRE in viral replication. Once the native CRE is inactivated, the relocated CRE becomes indispensable, and any deletion of the relocated CRE is expected to be lethal. Consequently, viruses retaining the insert and the relocated CRE are selectively favored, thereby enhancing reporter stability. This strategy proved effective in EV-A71. However, its effectiveness may vary across the species, as discussed below.

In HRV-B14, by contrast, CRE relocation did not stabilize the reporter insert as the reporter was lost by passage 5 in both the Mid_CRE and Mid_CRE/ΔCRE viruses (Fig. 5D). Notably, we infer that the deletions appeared to occur within mScarlet3 sequence while leaving the relocated CRE intact. As a result, the selective constraint that preserved the reporter insert in EV-A71 was not imposed in HRV-B14. Several factors may account for this species-specific difference. First, the position and sequence of the CRE vary among enteroviruses and rhinoviruses, overlapping different coding regions in poliovirus, CVB3, cardiovirus, HRV-A, and HRV-B (15–17, 27–30). These differences may influence how efficiently a relocated CRE functions in each viral background. Another possibility is that HRV-B14 has a more limited tolerance for insert length. Even with the optimized 2A cleavage site, both the Mid_CRE and Mid_CRE/ΔCRE viruses replicated less efficiently than rB14, with viral titers at 12 hpi falling below the detection limit (Fig. 5C), suggesting that insertion of the reporter gene had already imposed a substantial fitness cost. Addition of the 96-nt B14 CRE may therefore have increased the genetic burden beyond the tolerance of HRV-B14, favoring deletion within the mScarlet3 sequence while retaining the relocated CRE. Other factors, such as the structural stability or functional efficiency of the relocated CRE itself, may also contribute. For enterovirus species with limited tolerance for foreign insertions, combining CRE relocation with a shorter reporter gene may therefore be necessary to achieve stable reporter maintenance.

In summary, our modular strategy - (i) reporter insertion at the 5′UTR-VP4 junction region with optimization of the 2A protease cleavage site, (ii) GC optimization of the reporter gene, and (iii) CRE relocation – provides a general framework for constructing stable reporter enteroviruses and may be broadly applicable across the Enterovirus genus. A limitation of this study is that the strategy was experimentally validated in only two species, HRV-B and EV-A. Consequently, each component should be evaluated individually to determine the optimal design for a given enterovirus species.

In this study, we established a practical reporter HRV that enables direct visualization of viral infection and facilitates high-throughput quantitative analyses. By combining MidGC-B14 with airway epithelial organoids, we monitored the epithelial immune response over time while simultaneously tracking viral spread in real time (Fig. 7). The selective induction of type III interferons (IFN-*λ*) and of IP-10, a biomarker of HRV-induced asthma exacerbation, indicates that this system captures physiologically relevant features of the airway response to HRV. This combined system may therefore serve as a model for studying HRV-induced exacerbation of asthma and COPD. Furthermore, replacing mScarlet3 with a reporter better suited for deep-tissue imaging, such as a near-infrared fluorescent protein or a bioluminescent protein, together with the use of susceptible transgenic mouse models, could extend this approach to *in vivo* imaging, enabling noninvasive longitudinal monitoring of viral spread and tissue tropism at the whole-animal level.

More broadly, incorporating RNA structural analysis into the rational design of reporter viruses may improve insert stability not only in HRVs but also in other enteroviruses. Together, our findings provide both a practical platform for studying HRV biology and a general strategy for the development of genetically stable reporter viruses across the genus Enterovirus.

## MATERIALS AND METHODS

### Cells and viruses

H1-HeLa cells (ATCC, CRL-1958) were used for the propagation and titration of HRV-A1B and HRV-B14, and RD-S cells (RD cells stably expressing SCARB2) were used for EV-A71. RD-S cells were established as previously described (31). H1-HeLa cells were cultured in Eagle’s minimum essential medium (E-MEM) (Wako, 051-07615), and RD-S cells in Dulbecco’s modified Eagle medium (D-MEM) high glucose (Wako, 043-30085), each supplemented with 10% fetal bovine serum (FBS), 100 U/ml penicillin, and 100 µg/ml streptomycin, and both containing L-glutamine. Human airway epithelial cells derived from human iPSCs were manufactured according to established protocols as described before (20, 21) and provided by HiLung Inc. Briefly, lung progenitor cells were stepwise induced from human iPSCs following a 21-days and 4-steps protocol (20, 21). At day 21, lung progenitor cells were isolated with specific surface antigen carboxypeptidase M and seeded onto the upper chamber of a 24-well Cell Culture Insert (CORNING, 3470), followed by a 28-day differentiation of airway epithelial cells in air-liquid-interface culture. All cells were maintained at 37°C under 5% CO2. HRV-A1B (ATCC, VR-1645) and HRV-B14 (ATCC, VR-284) were obtained from the American Type Culture Collection. EV-A71 (strain SK-EV006) was generated by reverse genetics as previously described (9, 32). All infections were performed at 33°C.

### Plasmid construction

Four reverse genetics vectors were constructed, each combining a transcription promoter - RNA polymerase I (Pol I), Pol I with an upstream hammerhead ribozyme (P+hh), cytomegalovirus (CMV), or CMV with an upstream hammerhead ribozyme (C+hh) - with a downstream hepatitis delta virus (HDV) ribozyme and a transcription terminator. The human Pol I promoter, HDV ribozyme, and terminator cassette was synthesized (GenScript), with the Pol I promoter sequence based on a previous report (12). For the Pol I–based vectors, pUC19 (Nippon Gene, 311-01004) was digested with HindIII and EcoRI, and the promoter, HDV ribozyme, and terminator sequences were inserted. For the CMV-based vectors, pCI-neo (Promega, E1841) was digested with MluI and XmaI, and the HDV ribozyme was inserted. Viral genomes, or hammerhead ribozyme–viral genome cassettes, were then inserted into the Pol I vectors (digested with SwaI and NcoI; assembled with the NEBuilder HiFi DNA Assembly Master Mix, NEB, E2621S) or the CMV vectors (digested with BmtI and SmaI; assembled with In-Fusion Snap Assembly Master Mix, Takara, 638947).

The mScarlet3 sequence was amplified from pDx_mScarlet3 (a gift from Dorus Gadella; Addgene plasmid #189754; RRID:Addgene_189754) and inserted into the 5′UTR–VP4 region of the HRV-B14 genome together with a protease cleavage sequence. Six cleavage sequences were tested: the native 5-aa (wt5aa) and 6-aa (wt6aa) 2A sequences, the conserved SITTY and DSITTY 2A sequences, the native 3C sequence (wt3C), and a commercial 3C cleavage sequence (LEVLFQ). For HRV-B14, a fragment spanning part of the 5′UTR, mScarlet3, the cleavage sequence, VP4, and part of VP2 was amplified and inserted following digestion with SnaBI and StuI. The CRE was appended to the 3′ end of mScarlet3 using the same strategy. GC-content variants of mScarlet3, in which the native GC content (63%) was reduced to 38% across the entire gene (All) or within the 5′ (Pre), 3′ (Post), or central (Mid) region, were synthesized (GenScript). For CRE relocation, strains carrying inactivating mutations in the native-CRE (ΔCRE), a relocated CRE (Mid_CRE), or both (Mid_CRE/ΔCRE) were generated from the MidGC strain. Native-CRE inactivation was introduced using mutagenic primers spanning the CRE, with insertion via restriction sites flanking the CRE (common to HRV-B14 and EV-A71). All restriction enzymes were from NEB. Constructs were verified using a SeqStudio Genetic Analyzer (Thermo Fisher Scientific).

### Reverse genetics (virus recovery)

Viruses were recovered by transfecting genomic plasmids into the corresponding host cells. HRV genomic plasmids were transfected into H1-HeLa cells (0.5 µg/well in a 24-well plate) using FuGENE 4K (Promega, E5911), and EV-A71 genomic plasmids into RD-S cells (0.8 µg/well in a 24-well plate) using TransIT-LT1 (Mirus, MIR2304). For RNA transfection, genomic plasmids were linearized with BspQI, transcribed *in vitro* using the ScriptMAX Thermo T7 Transcription Kit (Toyobo, TSK-101), and the resulting RNA was introduced using TransIT-mRNA (Mirus, MIR2225). Culture supernatants and cells were harvested at 4 or 5 days post-transfection for HRV and for EV-A71, freeze-thawed once, clarified by centrifugation, and stored at –80°C.

### Plaque assay

Viral titers were determined by plaque assay on H1-HeLa cells (HRV) or RD-S cells (EV-A71) in 48-well plates. Cells were inoculated with 10-fold serially diluted virus and overlaid with an equal-volume mixture of 1.8% agar (Wako, 012-11872) and 2× medium prepared from 2× MEM (Thermo Fisher Scientific, 11935046) supplemented with FBS and penicillin–streptomycin. After incubation at 33°C for 3 days, cells were fixed by adding 4% paraformaldehyde to the overlay for 30 min, washed, and the agar was removed. Plaques were visualized by crystal violet staining (Nacalai Tesque, 09803-62). The same protocol was used for HRV and EV-A71.

### Alignment of 2A cleavage sequences

HRV-B genome sequences were retrieved from the NCBI GenBank nucleotide database using NCBI Entrez with the query txid147712[Organism] AND “complete genome”[Title] AND 7000:8000[SLEN]. For each genome, the VP1 and 2A regions were identified from the mat peptide annotations, and the corresponding amino acid sequences were obtained from the annotated translations or by translating the coding sequence. The cleavage-site sequence, defined as the six C-terminal residues of VP1 and the two N-terminal residues of 2A, was extracted for each strain. The resulting 32 sequences (Table S1) were aligned using MAFFT v7.526 with the auto option, and sequence conservation at the VP1–2A junction was visualized using WebLogo v3.9.0. The extraction script is available at [https://github.com/kazuma13/hrv-reporter-virus].

### Serial passage

Recovered viruses were serially passaged in H1-HeLa (HRV-B14) or RD-S (EV-A71) cells at an MOI of 0.01 at 33°C. The titer of P1 was determined, and the dilution required to achieve an MOI of 0.01 was applied for inoculation. Because each passage was harvested at the same stage, when complete cytopathic effect was observed, the same dilution scheme was used for subsequent passages without re-titration. Passaging was continued to obtain the fifth (P5) and tenth (P10) passages.

### Flow cytometry and reporter retention assay

Cells were infected with the indicated passage at an MOI of 0.1 and harvested at 12 hpi (HRV-B14, H1-HeLa) or 14 hpi (EV-A71, RD-S). Cells were fixed with 4% paraformaldehyde for 30 min at room temperature, washed with PBS, and permeabilized with 0.1% Triton X-100 for 30 min at room temperature (both Nacalai Tesque). For HRV, cells were stained with anti-VP3 (Invitrogen, MA5-18249; 1:100) followed by Alexa Fluor 488 goat anti-mouse IgG (Invitrogen, A-11001; 1:1,000); for EV-A71, with anti-3CD (GeneTex, GTX132355; 1:100) followed by Alexa Fluor 488 goat anti-rabbit IgG (Invitrogen, A-11008; 1:1,000). Primary antibodies were incubated at 4°C overnight and secondary antibodies at 37°C for 1 h. Fluorescence was acquired on an Attune NxT flow cytometer (Thermo Fisher Scientific). Compensation was applied for the green and red channels, and gating for positive cells was set against mock-infected controls. Reporter retention was calculated as the proportion of mScarlet3-positive cells among VP3-positive (HRV) or 3CD-positive (EV-A71) cells × 100. All assays were performed in at least three independent biological replicates.

### Immunofluorescence

Cells were infected at an MOI of 0.1 and incubated at 33°C for 12 h. Cells were fixed with 4% paraformaldehyde, permeabilized with 100% methanol for 2 min at room temperature, and stained with anti-VP3 (Invitrogen, MA5-18249; 1:100) followed by Alexa Fluor 488 goat anti-mouse IgG (Invitrogen, A-11001; 1:1,000). Nuclei were stained with DAPI (Dojindo, D523). mScarlet3 was detected by its intrinsic fluorescence. Mock-, rB14-, and Ori-B14-infected cells were compared. Images were acquired on a Nikon AX confocal microscope with NIS-Elements software.

### Growth curves

Cells were infected at an MOI of 0.01 (H1-HeLa for HRV-B14, RD-S for EV-A71). After adsorption for 60 min at 33°C with tilting every 15 min, cells were washed twice and supplied with growth medium. Culture supernatants were collected at 12, 24, 36, and 48 hpi, and viral titers were determined by plaque assay. Data are presented as mean ± SD.

### RT-PCR and sequencing

Viral RNA was extracted using the ISOSPIN Viral RNA kit (Nippon Gene). Reverse transcription was performed with SuperScript IV (Thermo Fisher Scientific) using both random hexamers and oligo(dT). PCR was performed with KOD One PCR Master Mix (Toyobo, KMM-201) according to the manufacturer’s protocol, using the following primers: HRV-B14, 5′-GGCCCCTGAATGCGGCTAACCTTAA-3′ (forward) and 5′-TTAGATGGGTCCATTGACAGTGATT-3′ (reverse); EV-A71, 5′-ATTGGCCATCCGGTGTGCAATAGAG-3′ (forward) and 5′-GTCTTGTTTGAGGCTCTGTTTGCCT-3′ (reverse). Amplicons were sequenced by Sanger sequencing using BigDye Terminator v3.1 on a SeqStudio Genetic Analyzer (Thermo Fisher Scientific).

### RNA structural analysis and GC content

The GC content of viral genomes was calculated using a custom Python script with a sliding window of 100 nt and a step of 50 nt. RNA secondary structures were predicted using RNAfold from the ViennaRNA Package v2.7.2 at 33°C. For Fig. 6, the junction region spanning the relocated CRE and ±100 nt of flanking sequence (the 3′ end of the Mid variant of mScarlet3, the CRE, and the 5′ end of VP4) was analyzed for seven strains: HRV-A16, HRV-B14, HRV-C15, EV-A71, CVB3, PV1, and EV-D68. All analysis scripts are available at [https://github.com/kazuma13/hrv-reporter-virus].

### Fluorescence imaging and immune response analysis in infected organoids

Airway organoids were infected with MidGC-B14 at an MOI of 0.1. At 0, 6, 12, 18, 24, 48, and 72 hpi, red fluorescence was imaged using a BZ-X800L microscope (Keyence). After confirming the presence of cells across the entire well by bright-field imaging, the red-fluorescent area was quantified using BZ-H4A analysis software (Keyence). Basolateral culture medium was collected at the same time points, and 13 interferons and cytokines were quantified using the LEGENDplex Hu Anti-Virus Response Panel 1 (13-plex) (BioLegend, 741269) on an Attune NxT flow cytometer (Thermo Fisher Scientific), according to the manufacturer’s instructions. Three independent wells were prepared for the mock group and four for the infected group. Concentrations below the LOD were substituted with the LOD value before log10 transformation. IFN-γ, IL-1β, IL-6, IL-12p70, and GM-CSF were below the LOD in all replicates of both groups at every time point and excluded from the statistical analysis.

### Statistical analysis

Statistical analyses were performed in R v4.6.1 using the base stats package, with data handling in dplyr v1.2.1 and tibble v3.3.1, and plotting in ggplot2 v4.0.3. Comparisons between two groups used Welch’s *t*-test; all-pairwise comparisons among more than two groups used the Games-Howell test; and comparisons against a single control used Welch’s *t*-test with Holm correction. Data are presented as mean ± SD for growth curves. All experiments were performed in at least three independent biological replicates. The statistical test used for each panel is indicated in the corresponding figure legend.

## ACKNOWLEDGMENTS

This study was supported by Japan Society for the Promotion of Science KAKENHI (21K17282), Japan Science and Technology Agency COI-NEXT (JPMJPF2018), the SENSHIN Medical Research Foundation, and Japan Science and Technology Agency Moonshot R&D (JPMJMS2025). These study sponsors had no role in the analysis, interpretation of data, writing of the report, and the decision to submit the paper for publication. During the preparation of this work, the authors used Claude (Anthropic) to assist with coding in R for figure generation and statistical analysis, and with English-language editing. After using this tool, the authors reviewed and edited the content as needed and take full responsibility for the content of the published article. We thank M. Tomiyama, C. Cheng, D. C. Patino, C. Kominami, M. Ishibashi, and K. Toyoda for their assistance.

## POTENTIAL CONFLICTS OF INTEREST

The authors declare that they have no conflicts of interest.

---

**FIG S1** Flow cytometry analysis of mScarlet3 expression for each 2A cleavage construct, related to Fig. 2C. H1-HeLa cells were transfected with genome plasmids encoding mScarlet3 with the indicated cleavage sequence (wt5aa, wt6aa, SITTY, DSITTY, wt3C, or LEVLFQ) between mScarlet3 and VP4, or with an empty vector lacking mScarlet3 (Empty), and harvested at 4 dpt. Histograms show mScarlet3 fluorescence intensity, and the gate for mScarlet3-positive cells (R2) was set using the Empty control. The percentage of mScarlet3-positive cells is indicated in each plot. Data are representative of three independent biological replicates.

**FIG S2** Flow cytometry analysis of reporter retention during serial passage, related to Fig. 4E and 4G. MidGC-B14, MidGC-EV-A71, and Ori-EV-A71 were serially passaged, and cells infected with the indicated passages were analyzed by flow cytometry. MidGC-B14 and MidGC-EV-A71 were analyzed at P1, P5, and P10, and Ori-EV-A71 at P1 and P5. For MidGC-B14, cells were stained with an anti-VP3 antibody, and mScarlet3 (y axis) and VP3 (x axis) signals are shown. For MidGC-EV-A71 and Ori-EV-A71, cells were stained with an anti-3CD antibody, and mScarlet3 (y axis) and 3CD (x axis) signals are shown. VP3- or 3CD-positive, mScarlet3-negative cells (lower right) indicate infection by a mScarlet3-deficient mutant. Data are representative of three independent biological replicates.

**FIG S3** Identification of an mScarlet3 region permissive for CRE insertion, related to Fig. 5. (A) Schematic of the three CRE insertion patterns within mScarlet3, in which the HRV-B14-derived CRE was inserted after the codon for amino acid 75 (75CRE), 186 (186CRE), or 229 (229CRE), compared with the original mScarlet3 (Original). (B) AlphaFold2-predicted structure of mScarlet3 carrying the CRE at each position. The original mScarlet3 is shown in gray, and the regions corresponding to the 75CRE (orange), 186CRE (green), and 229CRE (magenta) insertions are highlighted. (C) Genome plasmids encoding mScarlet3 with the CRE at each position, the original mScarlet3 (Original), or an empty vector (Empty), were transfected into H1-HeLa cells. To assess mScarlet3 expression, fluorescence was imaged (left), and the proportion of mScarlet3-positive cells, normalized to that of Original, was determined by flow cytometry at 24 h post-transfection (right). Data are presented as mean ± SD from three independent biological replicates. Statistical comparisons used the Games-Howell test (C); *P* values are indicated in each panel.

**FIG S4** Flow cytometry analysis of reporter retention during serial passage of CRE-relocated constructs, related to Fig. 5D and 5F. (A) Mid_CRE and Mid_CRE/ΔCRE of HRV-B14 were serially passaged, and cells infected with P1 and P5 were stained with an anti-VP3 antibody and analyzed by flow cytometry. mScarlet3 (y axis) and VP3 (x axis) signals are shown. (B) Mid_CRE and Mid_CRE/ΔCRE of EV-A71 were serially passaged, and cells infected with P1, P5, and P10 were stained with an anti-3CD antibody and analyzed by flow cytometry. mScarlet3 (y axis) and 3CD (x axis) signals are shown. VP3- or 3CD-positive, mScarlet3-negative cells (lower right) indicate infection by a mScarlet3-deficient mutant. Data are representative of three independent biological replicates.

## Notes

### Competing Interest Statement

The authors have declared no competing interest.

