## Supplementary Table 1 and figure S1-4 for "A versatile and stable reporter platform for human rhinovirus and other enteroviruses"

### Slide 1
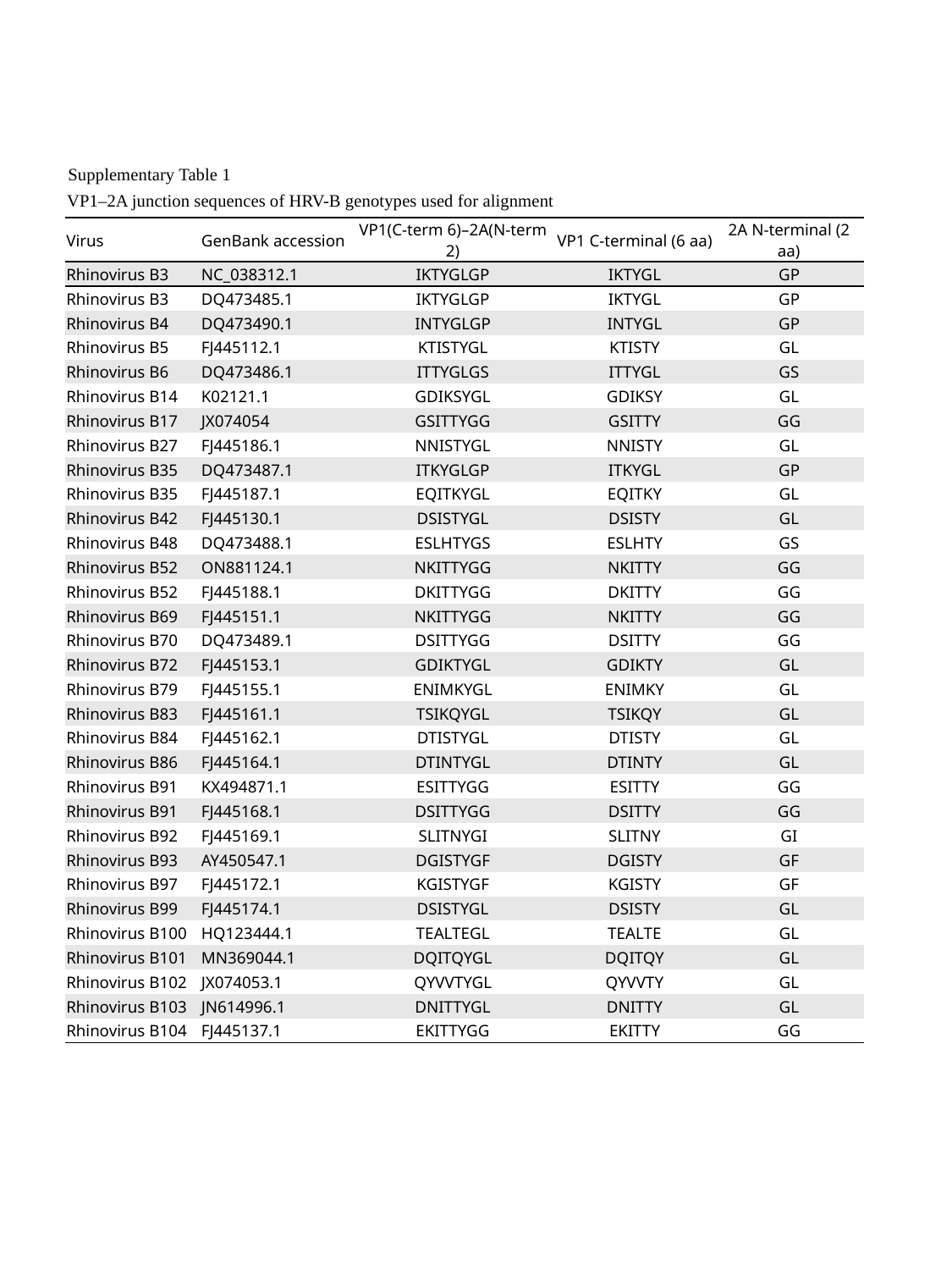

Supplementary Table 1
VP1–2A junction sequences of HRV-B genotypes used for alignment
| Virus | GenBank accession | VP1(C-term 6)–2A(N-term 2) | VP1 C-terminal (6 aa) | 2A N-terminal (2 aa) |
| --- | --- | --- | --- | --- |
| Rhinovirus B3 | NC\_038312.1 | IKTYGLGP | IKTYGL | GP |
| Rhinovirus B3 | DQ473485.1 | IKTYGLGP | IKTYGL | GP |
| Rhinovirus B4 | DQ473490.1 | INTYGLGP | INTYGL | GP |
| Rhinovirus B5 | FJ445112.1 | KTISTYGL | KTISTY | GL |
| Rhinovirus B6 | DQ473486.1 | ITTYGLGS | ITTYGL | GS |
| Rhinovirus B14 | K02121.1 | GDIKSYGL | GDIKSY | GL |
| Rhinovirus B17 | JX074054 | GSITTYGG | GSITTY | GG |
| Rhinovirus B27 | FJ445186.1 | NNISTYGL | NNISTY | GL |
| Rhinovirus B35 | DQ473487.1 | ITKYGLGP | ITKYGL | GP |
| Rhinovirus B35 | FJ445187.1 | EQITKYGL | EQITKY | GL |
| Rhinovirus B42 | FJ445130.1 | DSISTYGL | DSISTY | GL |
| Rhinovirus B48 | DQ473488.1 | ESLHTYGS | ESLHTY | GS |
| Rhinovirus B52 | ON881124.1 | NKITTYGG | NKITTY | GG |
| Rhinovirus B52 | FJ445188.1 | DKITTYGG | DKITTY | GG |
| Rhinovirus B69 | FJ445151.1 | NKITTYGG | NKITTY | GG |
| Rhinovirus B70 | DQ473489.1 | DSITTYGG | DSITTY | GG |
| Rhinovirus B72 | FJ445153.1 | GDIKTYGL | GDIKTY | GL |
| Rhinovirus B79 | FJ445155.1 | ENIMKYGL | ENIMKY | GL |
| Rhinovirus B83 | FJ445161.1 | TSIKQYGL | TSIKQY | GL |
| Rhinovirus B84 | FJ445162.1 | DTISTYGL | DTISTY | GL |
| Rhinovirus B86 | FJ445164.1 | DTINTYGL | DTINTY | GL |
| Rhinovirus B91 | KX494871.1 | ESITTYGG | ESITTY | GG |
| Rhinovirus B91 | FJ445168.1 | DSITTYGG | DSITTY | GG |
| Rhinovirus B92 | FJ445169.1 | SLITNYGI | SLITNY | GI |
| Rhinovirus B93 | AY450547.1 | DGISTYGF | DGISTY | GF |
| Rhinovirus B97 | FJ445172.1 | KGISTYGF | KGISTY | GF |
| Rhinovirus B99 | FJ445174.1 | DSISTYGL | DSISTY | GL |
| Rhinovirus B100 | HQ123444.1 | TEALTEGL | TEALTE | GL |
| Rhinovirus B101 | MN369044.1 | DQITQYGL | DQITQY | GL |
| Rhinovirus B102 | JX074053.1 | QYVVTYGL | QYVVTY | GL |
| Rhinovirus B103 | JN614996.1 | DNITTYGL | DNITTY | GL |
| Rhinovirus B104 | FJ445137.1 | EKITTYGG | EKITTY | GG |

### Slide 2
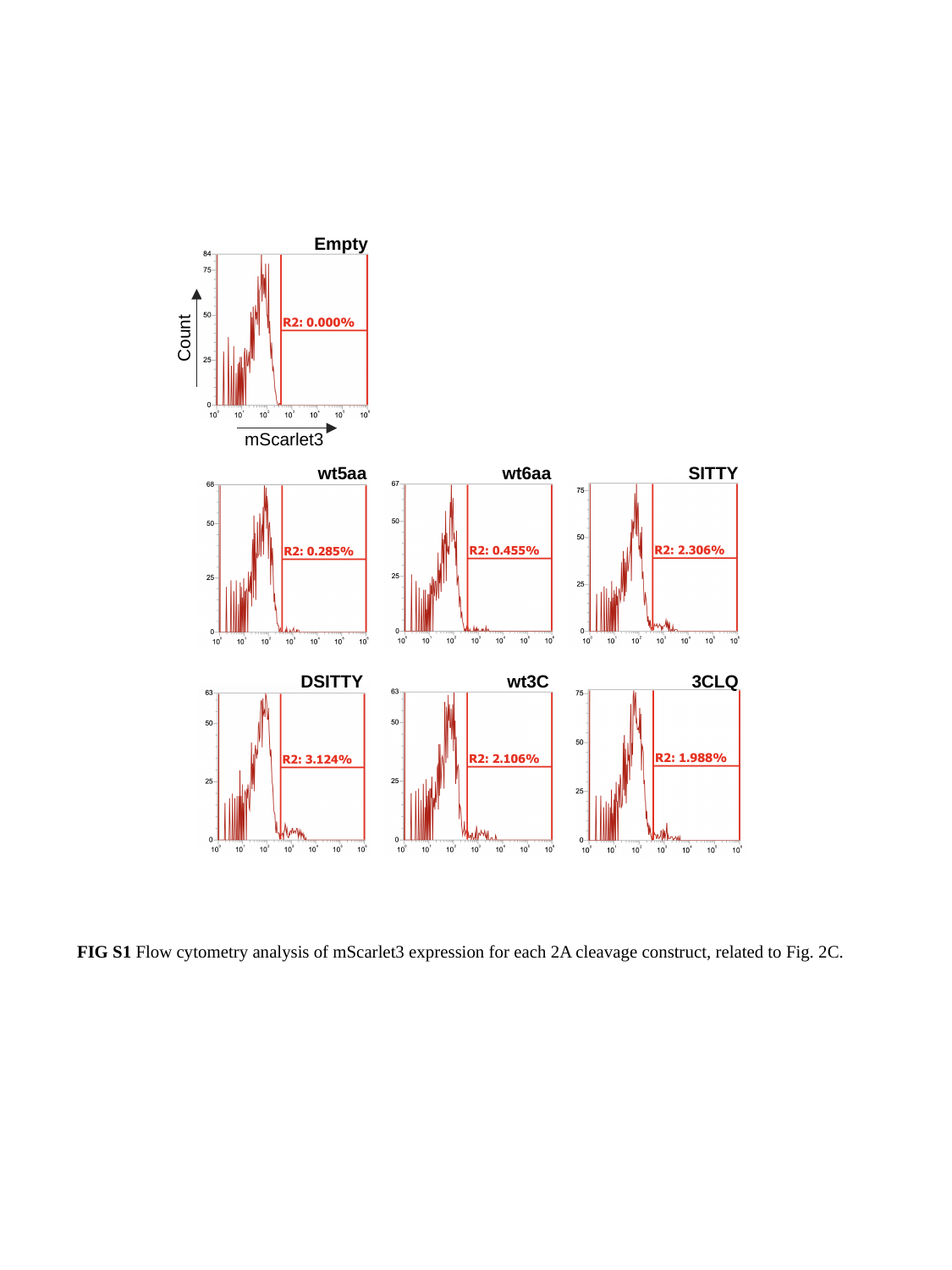

Empty
Count
mScarlet3
wt5aa
wt6aa
SITTY
DSITTY
wt3C
3CLQ
FIG S1 Flow cytometry analysis of mScarlet3 expression for each 2A cleavage construct, related to Fig. 2C.

### Slide 3
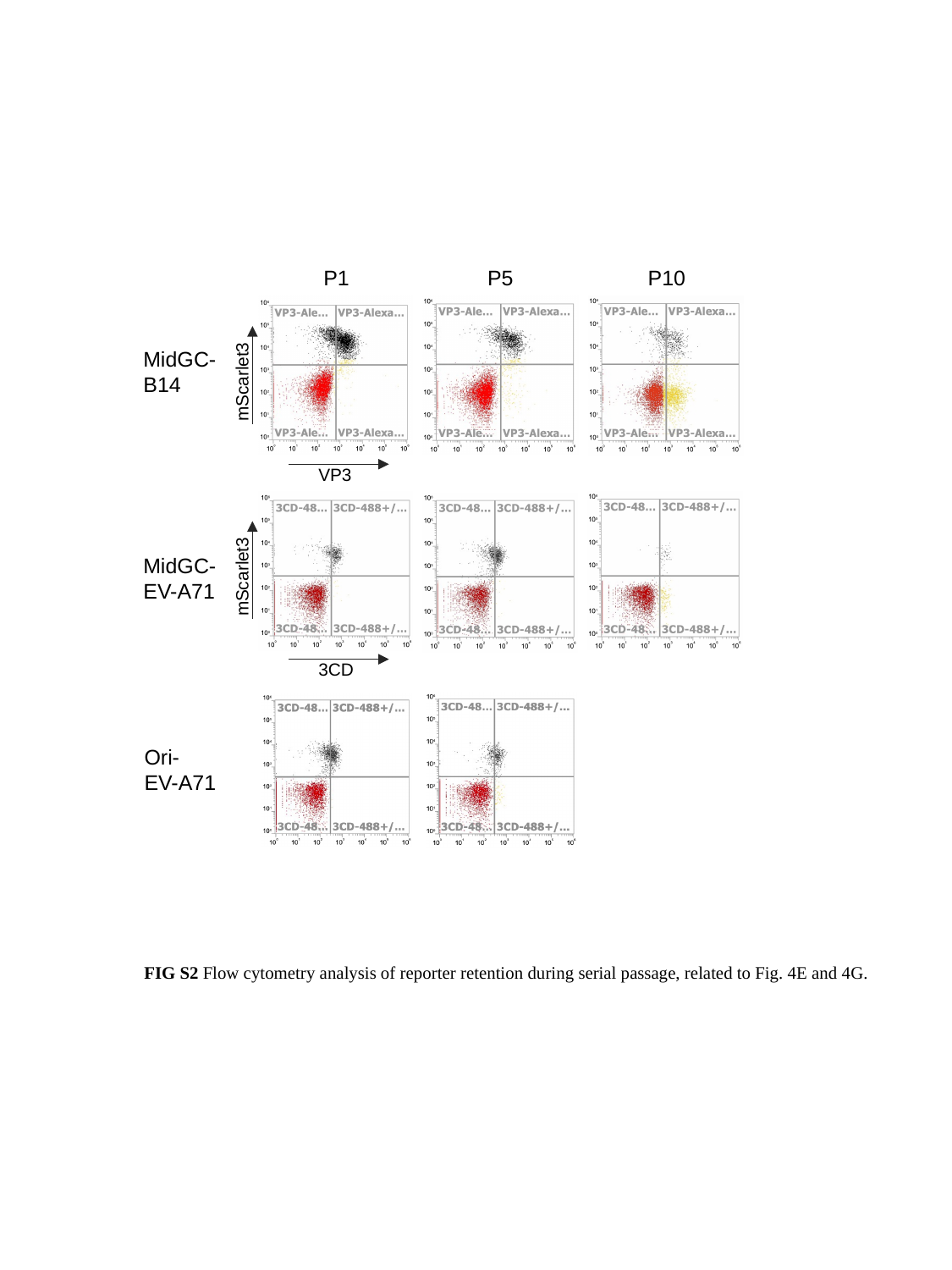

P1
P5
P10
MidGC-
B14
mScarlet3
VP3
MidGC-
EV-A71
mScarlet3
3CD
Ori-
EV-A71
FIG S2 Flow cytometry analysis of reporter retention during serial passage, related to Fig. 4E and 4G.

### Slide 4
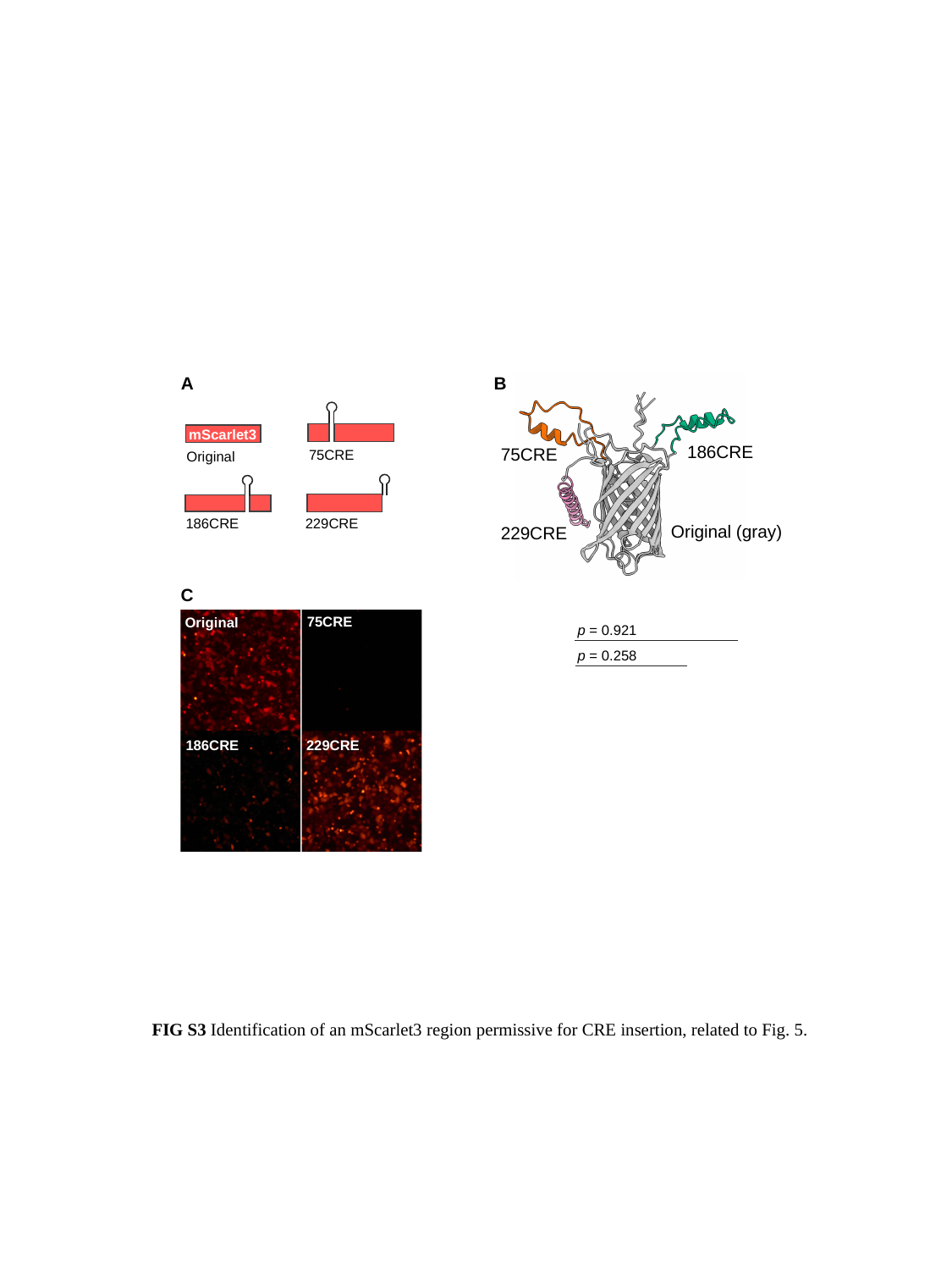

A
B
186CRE
75CRE
229CRE
mScarlet3
75CRE
Original
186CRE
229CRE
Original (gray)
C
75CRE
Original
p = 0.921
p = 0.258
186CRE
229CRE
FIG S3 Identification of an mScarlet3 region permissive for CRE insertion, related to Fig. 5.

### Slide 5
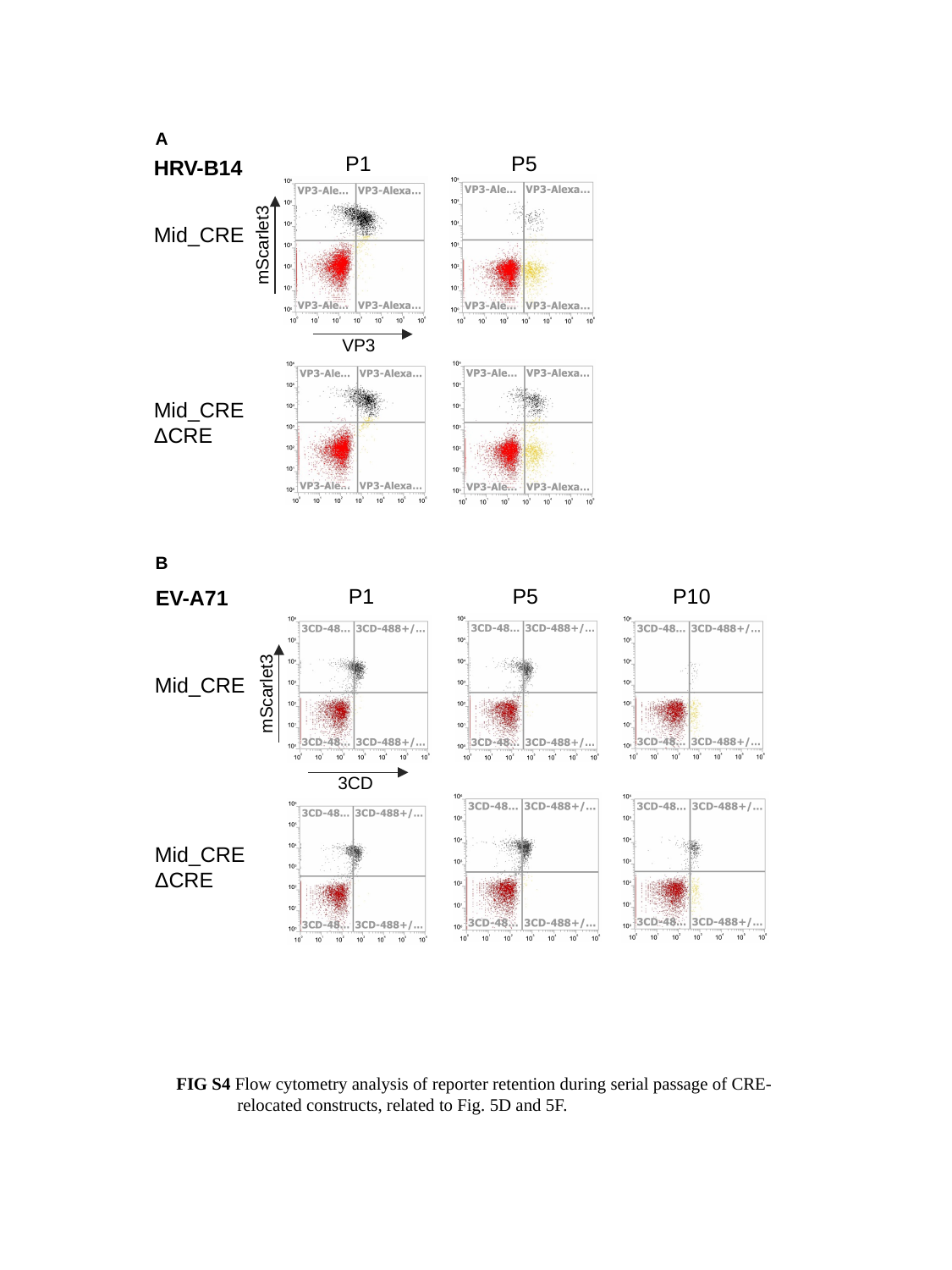

A
P1
P5
HRV-B14
Mid_CRE
mScarlet3
VP3
Mid_CRE
ΔCRE
B
P1
P5
P10
EV-A71
Mid_CRE
mScarlet3
3CD
Mid_CRE
ΔCRE
FIG S4 Flow cytometry analysis of reporter retention during serial passage of CRE-relocated constructs, related to Fig. 5D and 5F.
